# Resting state functional connectivity in relapsing remitting multiple sclerosis with mild disability – a data driven, whole brain multi-voxel pattern analysis study

**DOI:** 10.1101/2021.11.23.469578

**Authors:** Gowthami Nair, Sruthi S. Nair, K. M. Arun, Paul Camacho, Elshal Bava, Priya Ajayaghosh, Ramshekhar N. Menon, Muralidharan Nair, Chandrasekharan Kesavadas, Sheeba Arnold Anteraper

## Abstract

**Background:** Multivoxel pattern analysis (MVPA) has emerged as a powerful unbiased approach for generating seed regions of interest (ROIs) in resting-state functional connectivity (RSFC) analysis in a data-driven manner. Studies exploring RSFC in multiple sclerosis have produced diverse and often incongruent results.

**Objectives:** The aim of the present study was to investigate RSFC differences between persons with relapsing-remitting multiple sclerosis (RRMS) and healthy controls (HC).

**Methods:** We performed a whole-brain connectome-wide MVPA in 50 RRMS patients with expanded disability status scale ≤4 and 50 age and gender-matched HCs.

**Results:** Significant group differences were noted in RSFC in three clusters distributed in the following regions; anterior cingulate gyrus, right middle frontal gyrus, and frontal medial cortex. Whole-brain seed-to-voxel RSFC characterization of these clusters as seed ROIs revealed network-specific abnormalities, specifically in the anterior cingulate cortex and the default mode network.

**Conclusions:** The network-wide RSFC abnormalities we report agree with the previous findings in RRMS, the cognitive and clinical implications of which are discussed herein.

**IMPACT STATEMENT:** This study investigated resting state functional connectivity (RSFC) in relapsing remitting multiple sclerosis (RRMS) persons with mild disability (expanded disability status scale ≤4). Whole-brain connectome-wide multivoxel pattern analysis (MVPA) was used for assessing RSFC. Compared to healthy controls (HC), we were able to identify three regions of interest for significant differences in connectivity patterns, which were then extracted as a mask for whole-brain seed-to-voxel analysis. A reduced connectivity was noted in the RRMS group, particularly in the anterior cingulate cortex and the default mode network regions, providing insights into the RSFC abnormalities in RRMS.

## INTRODUCTION

Multiple sclerosis (MS) is a chronic inflammatory and progressive degenerative disease of the central nervous system (CNS). The hallmark feature of this disorder is the presence of disseminated lesions or ‘plaques’ in multiple areas within the brain, optic nerve and spinal cord (Noseworthy et al., 2000). These lesions are pathologically recognised as hubs of demyelination, inflammation, axonal loss and gliosis in the white and grey matter (Compston & Coles, 2008). The first symptomatic clinical episode of MS is referred to as clinically isolated syndrome (CIS) which can variously evolve into relapsing-remitting multiple sclerosis (RRMS) and secondary-progressive multiple sclerosis (SPMS) in the majority, while 10-15% have a progressive course from the onset (primary progressive multiple sclerosis: PPMS) (Lublin et al. 2014).

Technological advancements in neuroimaging have enhanced the diagnostic utility of magnetic resonance imaging (MRI) for MS (Fox et al., 2011). Beyond the common application of MRI for delineating structural abnormalities, functional MRI has emerged as a promising tool for understanding the anomalous brain networks in MS (Tahedl et al., 2018). Connectivity studies indicate that the connectivity or relay of information between different brain regions alters as the disease progresses (Tahedl et al., 2018). These alterations to the connectome impair the network functioning efficiency of brain (Liu et al., 2017; Shu et al., 2016). Abnormalities in resting state functional connectivity (RSFC) have been observed in all the subtypes of MS, including the very early stages. Synchronization in the RSFC components of the default mode network (DMN) was found to be higher in CIS compared to RRMS and healthy controls (HC). The compensatory cortical reorganization in the resting state networks which facilitate this diminishes in the later stages of the disease (Roosendaal et al., 2010). Cross-sectional studies have suggested a correlation for higher disability and poorer walking performance in MS with decreased RSFC in the default mode, visual, cognitive, and cerebellar networks (Bollaert et al., 2018; Rocca et al., 2018).

Connectivity studies in RRMS have shown a notable lack of consistency in their results which is attributable to the highly variable methodologies and the variability of clinical features of MS (Bonavita et al., 2011; Filippi et al., 2013; Sbardella et al., 2015). Using data driven techniques such as independent component analysis (ICA), it was noted that there was a significant reorganization of RSFC in networks involved in motor, sensory and cognitive processes between RRMS patients and HC (Faivre et al., 2012). Another study noted that the DMN had weaker RSFC in the anterior and central posterior cingulate cortices in RRMS patients, where the cognitively preserved patients had more significant variations than cognitively impaired patients (Bonavita et al., 2011). More recent studies in RRMS have generated similar results with reduced connectivity demonstrated in networks like the default mode, attention and memory networks (Nejad-Davarani et al., 2016).

Analysis of resting state functional MRI data follows a hypothesis-driven or a data-driven methodology. In the hypothesis-driven approach, the investigator predefines the regions of interest (ROIs) or seeds which are usually guided by results of previous studies or other hypotheses. Techniques such as seed-to-voxel, seed-to-seed, ROI-to-ROI, and higher-level graph theory analysis have been utilised to study the underlying network abnormalities in persons with MS (Tahedl et al., 2018). This approach has an inherent risk of bias as the selection of the predefined regions is subjective.

The data-driven whole brain approach is more flexible and open without any pre-set notions. This method helps in finding new components and provides different insights into existing problems (Iraji et al., 2016). A widely used approach in MS is ICA. ICA helps to detect underlying patterns in the functional MRI (fMRI) signal without any prior knowledge about the task or shape of the hemodynamic response. It separates components in the data to identify spatially independent brain networks. Each non-noise component represents a functional network over a time course thereby allowing identification of independent signals from a combination of signals (McKeown et al., 1998; Tahedl et al., 2018). Multivoxel pattern analysis (MVPA) is another data-driven approach to study RSFC. MVPA attempts to discover spatial patterns of fMRI activity that correspond to different experimental conditions (Mahmoudi et al., 2012). It identifies distributed response patterns across voxels within a given time course. In this approach, seed-points are extracted from the group for further whole-brain connectivity analysis (Mahmoudi et al. 2012). MVPA has been sparingly used but is a potentially useful tool for studying RSFC in MS, which is a disease characterized by spatially heterogeneous damage and adaptations to neural pathways (Dineen et al. 2009, Sbardella et al. 2015). The objective of the present study was to examine the RSFC variations in RRMS persons with mild physical disability compared to HCs using MVPA technique.

## METHODS

### Ethics

The study commenced after approval from the Institutional Ethics Committee. All participants provided written informed consents prior to recruitment.

### Participants

The study was conducted as part of a larger project on cognitive dysfunction and multimodality MRI in RRMS. For this study, we recruited 50 persons with RRMS aged 18 years or above from the Multiple Sclerosis Clinic of the Institute who fulfilled the criteria for RRMS based on the 2010 revisions of the McDonald criteria (Polman et al., 2011) and were rated at or below 4 on the Kurtzke expanded disability status scale (EDSS). Participants were recruited from December 2017 to June 2020. Potential participants were excluded based on diagnosis of progressive forms of MS, disease onset before 12 years of age, recent clinical relapse, any corticosteroid therapy within one month, or significant systemic co-morbidities. At recruitment, a detailed clinical assessment was done to record the disease details, neuropsychological scores, functional scores (FS) and EDSS. Fifty age- and gender-matched HCs were also recruited for the study.

### Data acquisition

The MRI acquisition was performed using the 3T MRI (GE Discovery MR750w) with its standard 24 channel head array coils. Structural images were acquired first with T1-weighted imaging: TR = 12.6 ms, TE = 5.8 ms, time of inversion = 550 ms, flip angle = 12°, slice thickness = 2 mm, volumes = 1, slices = 156, imaging matrix = 512 × 512, and in-plane resolution = 0.35 × 0.35 mm^2^. The resting state fMRI sequences were acquired next with the participants instructed to keep their eyes open and stay awake. The protocol used was TR = 3000 ms, TE = 30 ms, flip angle = 80°, slice thickness = 3.2 mm, volumes = 150, imaging matrix = 64 × 64, and in-plane resolution = 3.31 × 3.31 mm^2^.

### Data pre-processing

Pre-processing and RSFC analysis were performed using the CONN toolbox (Whitfield-Gabrieli & Nieto-Castanon, 2012). The DICOM images were converted into the Neuroimaging Informatics Technology Initiative (NIfTI) format for further processing. Pre-processing consisted of removal of the initial five fMRI volumes, followed by functional realignment and unwarping based on field maps. These images then underwent slice-timing correction, followed by functional segmentation and structural segmentation of the T1-weighted images into grey matter, white matter and cerebrospinal fluid. The segmented images were then normalized onto the Montreal Neurological Institute (MNI) template space. This was followed by functional normalization and outlier detection (0.5 mm scan-to-scan motion). Finally, the images were smoothed with a full width half maximum (FWHM) Gaussian kernel of 8 mm. Temporal band-pass filtering was performed for a frequency range between 0.008 to 0.09 Hz.

### Multivoxel pattern analysis

Whole-brain MVPA identified seed ROIs for further characterization. The MVPA employs a voxel-to-voxel approach that generates ROIs for standard seed-to-voxel analysis. First, a principal components analysis (PCA) was used to reduce the dimensionality of the resultant data by estimating a multivariate representation of the connectivity pattern by computing the pairwise connectivity pattern between each voxel and the rest of the brain. A second PCA was run next across all participants to retain a conservative 25:1 ratio of subjects-to-components (4 in this study).

### Second level analysis

An omnibus F-test was performed on all 4 group-MVPA components simultaneously to identify the voxels that show significant differences in connectivity patterns between the groups/contrast of interest. We used a height threshold of p < 0.001 and a cluster-size false discovery rate (FDR) correction of 0.05 (non-parametric statistics). This resulted in three clusters.

### Post-hoc characterization

The three regions which were identified by MVPA were used as seeds to perform a whole-brain seed-to-voxel RSFC characterization. A one sample t-test was utilised separately for all three seeds on the HC and RRMS groups at a height-threshold of p < 0.001 and cluster threshold of p < 0.05 FDR-corrected.

## RESULTS

### Participants

The study had 32 females and 18 males each in the RRMS and HC groups. The mean age for the RRMS group was 32.8 (± 8.0) years and the HC group was 31.96 (± 7.5) years. The clinical characteristics of the participants are described in Table 1.

**Table 1:** Demographic and clinical characteristics of the study participants

|  | **Persons with**<br>**RRMS**<br>**n = 50** | **Healthy**<br>**controls**<br>**n = 50** | **P value** |
| --- | --- | --- | --- |
| **Demographic parameters** |  |  |  |
| Age(years); mean (SD) | 32.80 (8.03) | 31.96 (7.52) | 0.591 |
| Females; n (%) | 32 (64) | 32 (64) | - |
| Years of formal education: mean (SD) | 15.16 (2.17) | 16.1 (1.6) | 0.015 |
| **Clinical parameters** |  |  |  |
| Age of symptom onset (years);<br>mean (SD) | 26.78 (7.35) | - |  |
| Duration of illness (years);<br>mean (SD) | 6.15 (4.66) | - |  |
| Disease modifying therapy; n (%) | 43 (86) | - |  |
| EDSS; median (IQR) | 2 (1 – 2.5) | - |  |
| **Neuropsychology test scores** |  |  |  |
| ACE III-R; mean (SD) | 90.6 (6.0) | 96.4 (2.9) | <0 .001 |
| MMSE; mean (SD) | 29.0 (1.6) | 29.9 (0.303) | <0.001 |

Abbreviations: ACE III-R, Addenbrooke's Cognitive Examination III - Revised; EDSS, Expanded Disability Status Scale; HC, Healthy Control; IQR, Interquartile range; MMSE, Mini-mental state examination; n, number; RRMS, Relapsing Remitting Multiple Sclerosis; SD, standard deviation.

### Group MVPA results

After executing a whole-brain connectome-wide MVPA between the two groups, we found three statistically significant clusters distributed over anterior cingulate gyrus (ACC), right middle frontal gyrus (MFG) and the frontal medial cortex (Figure 1, Table 2).

**Figure 1:**
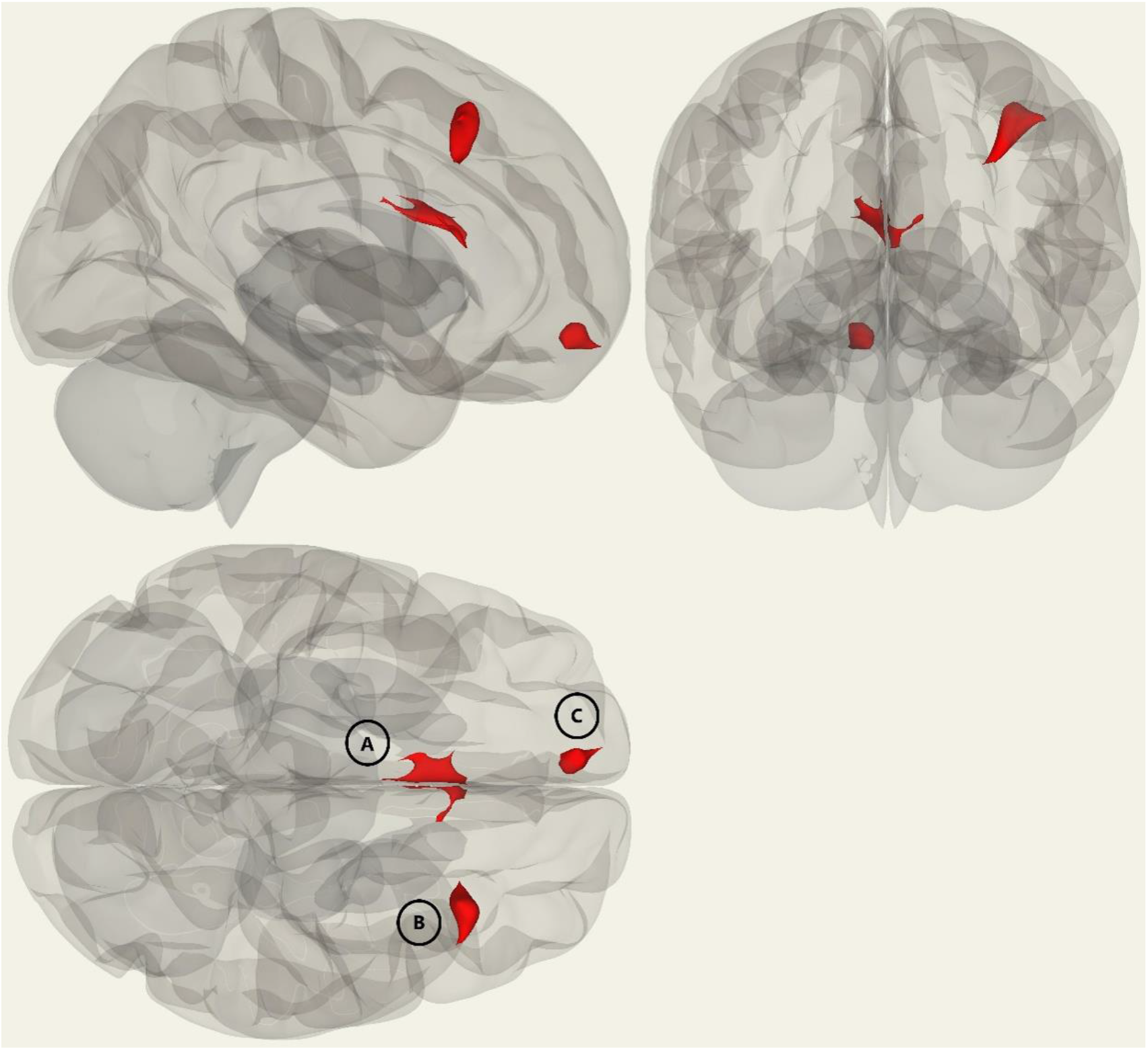
Whole-brain MVPA results indicating the regions that show significant differences in connectivity patterns when comparing the two groups (RRMS vs. HC). Cluster size (k) threshold of k > 99 (p < 0.05 FDR-corrected) and height-threshold of p < 0.001 (uncorrected, non-parametric statistics) were used to derive seed regions of interest: (A) anterior cingulate gyrus, (B) right middle frontal gyrus, and (C) frontal medial cortex.

**Table 2:** Whole-brain MVPA results indicating the regions that show significant differences in connectivity patterns when comparing the two groups (RRMS vs. HC).

| Brain regions | Peak cluster Coordinates | Voxels per cluster |
| --- | --- | --- |
| Anterior cingulate gyrus | -8 +12 +24 | 181 |
| Right middle frontal gyrus | +34 +24 +46 | 173 |
| Frontal medial cortex | -8 +56 - 16 | 99 |

MVPA: multi-voxel pattern analysis. A height threshold of whole-brain $p < 0.001$ and FDR-corrected cluster threshold of $p < 0.05$ (nonparametric statistics) was used to generate seed regions of interest.

### Post-hoc seed-to-voxel analysis of MVPA results

The MVPA cluster one was identified as ACC. Marked differences were noted between the two groups with this region as seed. In the RRMS group, the majority of the voxels clustered in the anterior cingulate gyrus and displayed an increased connectivity. A decreased connectivity was noted only in the left MFG. A wider area of variable connectivity was noted in HC with ACC as the seed. Like the RRMS group, ACC covered the greatest number of voxels with an increased connectivity in HC. In addition, ACC showed a positive connectivity in the right frontal pole, bilateral insular cortices, left anterior supramarginal gyrus (SMG), left MFG, right posterior SMG, right superior frontal gyrus (SFG) and bilateral frontal orbital cortices (Figure 2). Notably, left MFG had increased connectivity in HC in contrast to the RRMS group. A decrease in the connectivity was seen in HC over regions covering the right intracalcarine cortex, left cuneal cortex, bilateral lingual gyrus, vermis 8, right caudate and left hippocampus.

**Figure 2:**
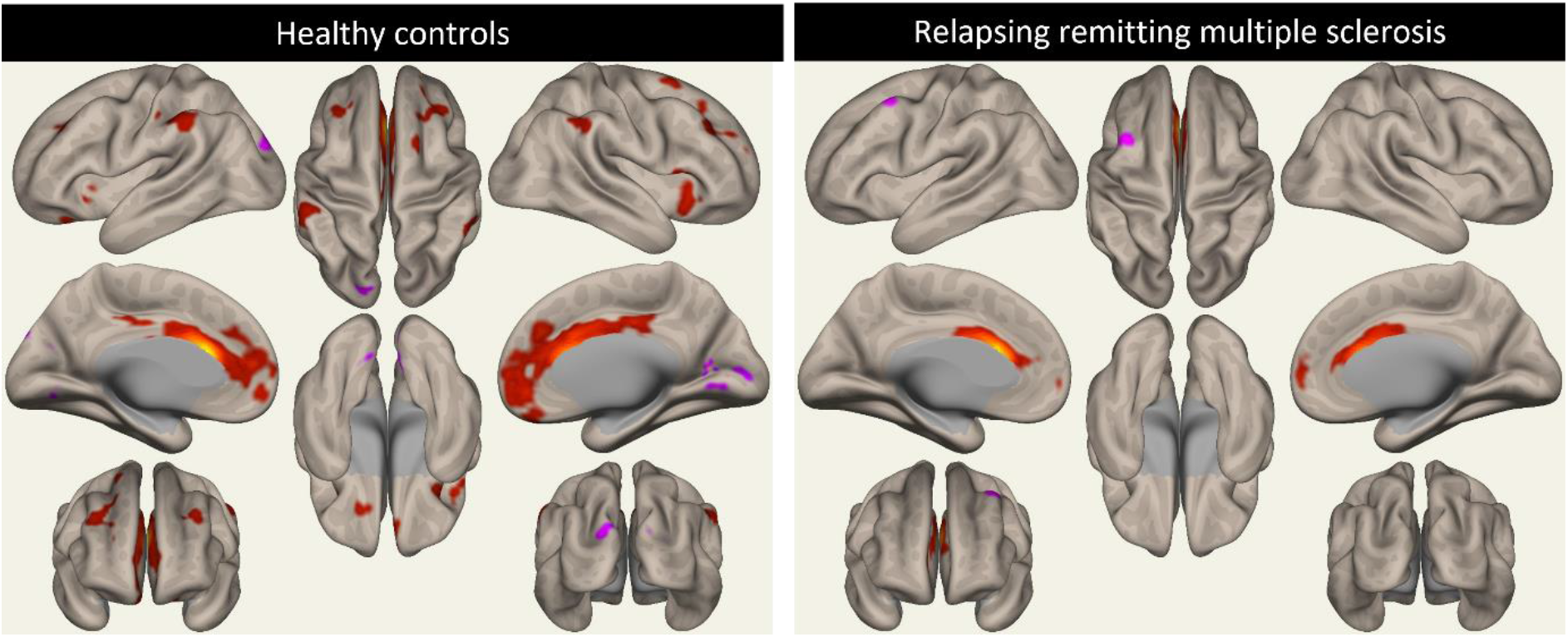
Results from second-level seed-to-voxel RSFC analysis for RRMS and HC groups (one-sample t-tests with height -threshold of p<0.001 and cluster-threshold of p<0.05 (FDR-corrected)) for MVPA-derived cluster A (anterior cingulate gyrus).

The second cluster was identified as the right MFG and on seed-to-voxel analysis showed an overlap in connectivity in nearly all the regions in both HC and RRMS (Figure 3). In both the groups the positive contrast was seen in the right and left frontal pole, right and left LOC, left cerebellum crus 2, left and right MTG and left cerebellum 9. Along with these regions, in the RRMS group right cerebellum crus2 showed a positive connectivity and, in the HC group, the precuneus and right inferior temporal gyrus had increased connectivity. A negative connectivity was noted in the right precentral gyrus, left occipital pole and ACC in both the groups. In the RRMS group, there was decreased connectivity in the right insular cortex, left frontal pole, left cerebellum 8, left precentral gyrus, right lingual gyrus, right occipital pole, right caudate, the brainstem, left hippocampus and left cerebellum 6. Overall, the clusters in the RRMS group were larger compared to the HC group.

**Figure 3:**
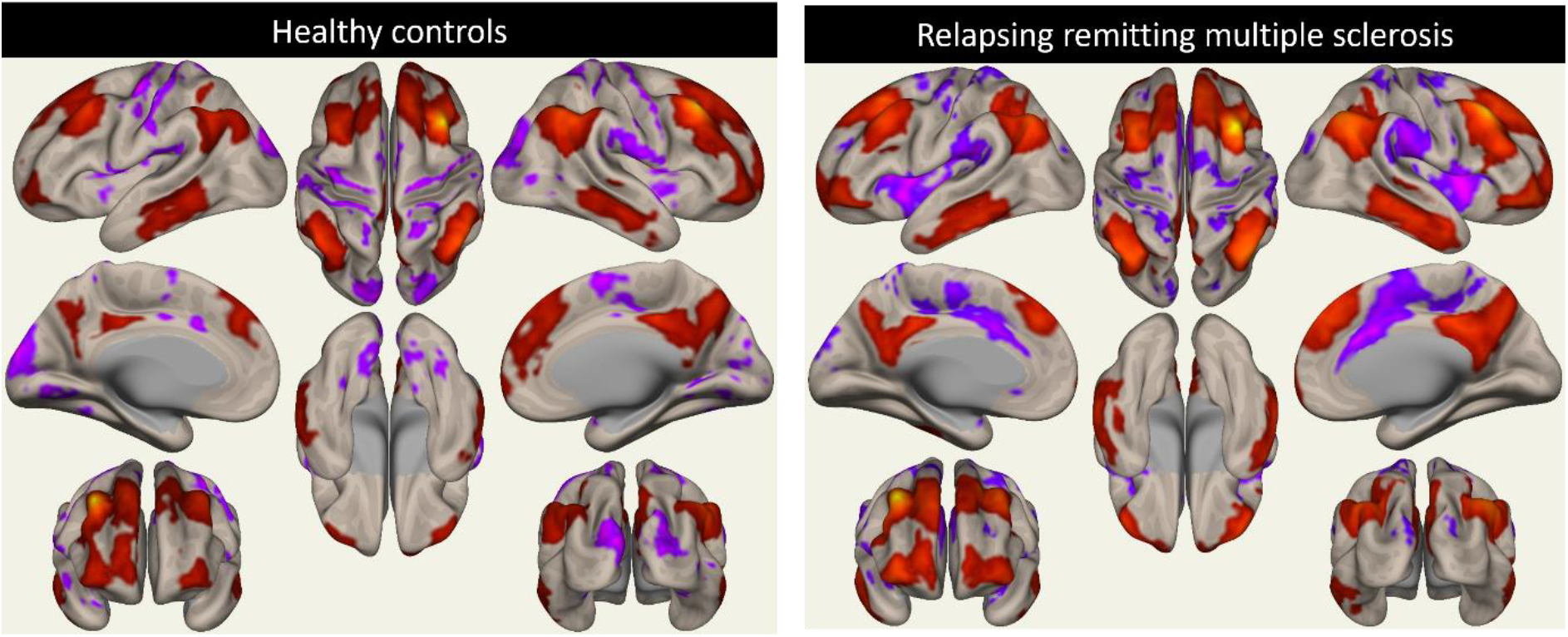
Results from second-level seed-to-voxel RSFC analysis for RRMS and HC groups (one-sample t-tests with height -threshold of p<0.001 and cluster-threshold of p<0.05 (FDR-corrected)) for MVPA-derived cluster B (right middle frontal gyrus).

The third cluster was located at the frontal medial cortex, also known as the medial prefrontal cortex (mPFC). When placed as the seed region, significant anti-correlation was seen between the RRMS and HC groups (Figure 4). RRMS group had increased connectivity in the right frontal pole, posterior cingulate gyrus, left superior frontal gyrus, right frontal pole, right and left superior LOC and right MTG and decreased connectivity in the right frontal operculum, left supplementary motor area, brainstem and right cerebellum crus 2. In the HC group, there was an increased activation in the right frontal pole, precuneus, right temporal pole, left and right LOC, right parahippocampal gyrus and left cerebellum 9 while connectivity was decreased in the left superior parietal lobule, right superior LOC, right inferior frontal gyrus, left insular cortex, left and right superior frontal gyri, right and left frontal poles, left inferior temporal gyrus, right and left cerebellum 8, left and right cerebellum 6, right middle temporal gyrus, left occipital pole and left lingual gyrus.

**Figure 4:**
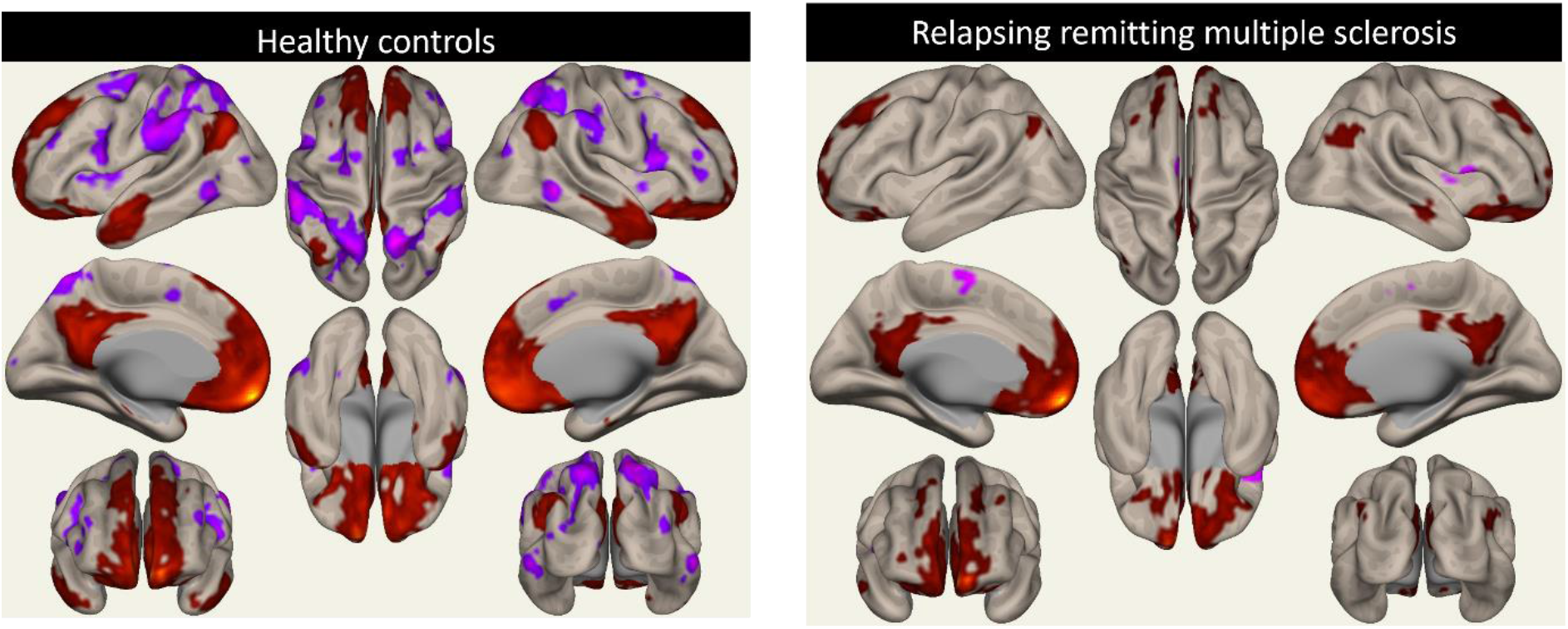
Results from second-level seed-to-voxel RSFC analysis for RRMS and HC groups (one-sample t-tests with height -threshold of p<0.001 and cluster-threshold of p<0.05 (FDR-corrected)) for MVPA-derived cluster C (frontal medial cortex).

## DISCUSSION

The main objective of this study was to investigate RSFC differences between RRMS and HC. We identified the following 3 significant clusters of altered connectivity in group MVPA: (1) ACC (2) right MFG and (3) mPFC. These clusters correspond to regions in various networks associated with language, attention, executive functions and emotional expression (Euston et al., 2012; Koyama et al., 2017; Semendeferi et al., 2001; Stevens et al., 2011; Jobson et al., 2021). The general pattern revealed in the post-hoc analysis was that of decreased RSFC in two of the three clusters in the RRMS group. Marked variability could be demonstrated in the connectivity patterns in RRMS and control groups in post-hoc analysis, particularly in anterior cingulate gyrus and DMN. These results could reflect the specific cognitive and higher order motor and sensory deficits in MS or the compensatory reorganization of networks congruous to the disease stage and disability.

The first cluster, ACC, has been credited with a major role in higher executive functioning and is an integral part of the dorsal attention network (DAN) (Loitfelder et al., 2012). The DAN is active during cognitive tasks pertaining to attention and mental control and is considered vital in the process of neuro-aging. Additionally, the ventral ACC has been associated with the anterior parts of the DMN. (Cera et al., 2019). ACC has also been implicated in integrating emotional and sensory information (Downar et al., 2000; Menon, 2015). Dorsal ACC is part of the salience network, which plays a crucial role in identifying relevant stimuli from our immediate surroundings to help guide behaviour (Menon, 2015; Qadir et al., 2018). Functionally, ACC has been linked to decision making, social interactions, emotional responses, and control of motor responses (Lavin et al., 2013; Menon, 2015).

A clinical study in RRMS found that changes in the connectivity patterns of the ACC were indicative of the restoration of attention and executive functions in the patients (Parisi et al., 2014). Additionally, cortical thinning of the ACC in MS has been linked to specific lateralized deficits; impaired verbal fluency and figural fluency were associated with left and right ACC thinning respectively (Geisseler et al., 2015). Another study looked at the correlation of RSFC in RRMS persons who underwent cognitive rehabilitation in information processing and executive function domains. After a 12-week rehabilitation, the intervention group displayed an increased RSFC between ACC and the right MFG compared to the baseline (Parisi et al., 2014). In our study, left MFG had increased connectivity in the HC group but had a decreased FC in the RRMS. Decreased activation of left MFG was registered along with abnormalities in DMN activation in MS compared to controls using a working memory network recruiting fMRI paradigm (Vacchi et al., 2016). Also, a general comparison between the connectivity maps (figure 2 & 3) shows the degree of under-connectivity present in the RRMS. Broadly, our findings agree with those identified from previously published studies.

The second cluster, right MFG, is associated with both the dorsal and ventral attention networks and acts by reorienting and interrupting attention to novel stimulus (Corbetta et al., 2008, Fox et al., 2011). The MFG has also been described as being involved in cognitive functions involving literacy and numeracy (Koyama et al., 2017) and as a core region in attention (Japee et al., 2015). When comparing the connectivity maps for each of the groups individually, overall similar connectivity patterns over the same regions can be delineated. However, the HC group had increased FC to the precuneus and the inferior temporal gyrus. A recent study in sleep-deprived healthy adults noted that disruption in the connectivity between the right precuneus and right MFG was correlated with decline in attention. The connectivity between these two regions were found to be an important node to maintain the function of attention (Li et al., 2020). These latter results suggest that focal decline in RSFC may correlate with cognitive dysfunction in RRMS.

Decreased RSFC in the RRMS group was seen in the right insular cortex. A recent study identified a motivational fatigue network consisting of the dorsal ACC, dorsolateral prefrontal cortex (PFC), ventromedial PFC and insula. This network monitors the internal state of the body and makes decisions regarding the costs and benefits associated with perseverating on a task and was demonstrated to be less efficient in MS persons (Chen et al., 2020). Our results could also support the framework of cognitive fatigue noticed in MS patients. Also, MFG registered a decreased RSFC to insular cortex and ACC which are components of the salience network (Menon, 2015). Previous studies have linked depression in MS with abnormal RSFC in important networks including DMN and salience networks. Within the salience network, an increased connectivity to right MFG was noted in depressed MS persons compared to non-depressed counterparts (Bonavita et al., 2016).

Using the mPFC as a seed region elucidated strikingly different patterns of RSFC in HC and RRMS. This region is part of the DMN and has been linked to multiple cognitive functions including decision making and action monitoring (Moreira et al., 2016). Indeed, decision-making time was correlated with diffusion abnormalities in mPFC, mFG and ACC in MS compared to HC (Mulhert et al. 2014). Aberrant medial frontal activation resulted from administration of tasks with increasing complexity in RRMS suggesting adaptive recruitment of this region to maintain cognitive performance (Bonnet 2010). The regulatory role of the mPFC has also been highlighted for functions such as attention, inhibitory control, habit formation and working, spatial or long-term memory. Additionally, this region has been identified in having several interconnections with subcortical regions such as the thalamus, amygdala and hippocampus (Jobson et al., 2021). A recent study found that the dynamic functional connectivity in specific regions such as the mPFC and precuneus could have a role in fatigue that is independent of physical disability (Tijhuis et al., 2021). So, in the RRMS patients, though early tests may fail to capture specific physical signs, their overall feeling of tiredness and poor exercise tolerance may in part be explained by faulty connectivity in these regions. The increase in connectivity between the mPFC and frontal pole has also been associated with facilitation of stable performance on complex speed-dependant information processing (Wojtowicz et al., 2014). These anti-correlated regions with mPFC were strikingly absent in the RRMS group, speaking to the slowly progressive cognitive decline seen in RRMS patients.

The primary strength of our study was the use of a data-driven approach based on MVPA which allowed for an unbiased method for comparison of RSFC between persons with RRMS and age- and sex-matched HC. This is one of the pioneer studies where this technique has been used in MS. An important limitation was that we did not analyse the information on the correlation of neuropsychological test scores and physical disability with RSFC abnormalities within our sample. This data could have provided more directly translatable clinical insight into the RSFC changes in each network with respect to disability in RRMS. Similarly, we did not correlate the network changes with structural imaging studies of cortical thickness or diffusion tensor imaging. The cross-sectional nature of the study precluded hypotheses on causal nature of the observed changes and information on network dysfunction versus compensatory changes in connectivity. The use of a relatively homogeneous group of RRMS with mild disability was considered important as the resting state data in MS tends to show marked variation over the disease course (Sbardella et al. 2015). However, even this group included patients with a notable variability in duration, symptoms, and MRI lesion load. The duration of formal education was not comparable between the RRMS and HC groups and the influence of the same in network connectivity is not well-understood. Future studies focussing on structural MRI and clinical correlation with the networks affected could help in better understanding of the mechanisms of cognitive dysfunction in RRMS. Long term longitudinal studies are essential to identify the moulding of networks in patients over time and their correlation with the disease course and disability in MS.

We have reported the RSFC abnormalities in patients of RRMS using data-driven MVPA. The post-hoc results revealed a general pattern of reduced RSFC in RRMS persons, particularly in the ACC and DMN. Precuneus and lateral occipital cortex were seen to have consistent abnormalities with decreased connectivity in the HC group. Previous studies of RSFC in RRMS have shown wide variability in the techniques and outcomes (Bonavita et al., 2011; Filippi et al., 2013; Sbardella et al., 2015). Further research utilising MVPA could provide unbiased imaging biomarkers for monitoring sub-clinical disease progression and intervention response in RRMS. MVPA-based RSFC abnormalities may be useful in understanding the mechanisms that alter the networks in RRMS and their influence on the highly variable clinical presentations observed across patients.

## AUTHOR CONTRIBUTION STATEMENT

| Author | Contribution |
| --- | --- |
| Gowthami Nair | Data collection, analysis and preparation of the manuscript and figures |
| Sruthi S. Nair | Conception of the study, data collection, preparation and critical revision of the manuscript |
| K. M. Arun | Data analysis and interpretation and final approval of manuscript |
| Paul Camacho | Critical revision of the manuscript |
| Elshal Bava | Data collection and analysis |
| Priya Ajayaghosh | Data collection |
| Ramshekhar N. Menon | Critical revision and final approval of the manuscript |
| Muralidharan Nair | Conception of the study and final approval of the manuscript |
| Chandrasekharan Kesavadas | Conception of the study, data analysis, and critical revision of the manuscript |
| Sheeba Arnold Anteraper | Data analysis and interpretation, critical revision and final approval of the manuscript |

